# Multiple early introductions of SARS-CoV-2 into a global travel hub in the Middle East

**DOI:** 10.1101/2020.05.06.080606

**Authors:** Ahmad Abou Tayoun, Tom Loney, Hamda Khansaheb, Sathishkumar Ramaswamy, Divinlal Harilal, Zulfa Omar Deesi, Rupa Murthy Varghese, Hanan Al Suwaidi, Abdulmajeed Alkhajeh, Laila Mohamed AlDabal, Mohammed Uddin, Rifat Hamoudi, Rabih Halwani, Abiola Senok, Qutayba Hamid, Norbert Nowotny, Alawi Alsheikh-Ali

## Abstract

International travel played a significant role in the early global spread of SARS-CoV-2. Understanding transmission patterns from different regions of the world will further inform global dynamics of the pandemic. Using data from Dubai in the United Arab Emirates (UAE), a major international travel hub in the Middle East, we establish SARS-CoV-2 full genome sequences from the index and early COVID-19 patients in the UAE. The genome sequences are analysed in the context of virus introductions, chain of transmissions, and possible links to earlier strains from other regions of the world. Phylogenetic analysis showed multiple spatiotemporal introductions of SARS-CoV-2 into the UAE from Asia, Europe, and the Middle East during the early phase of the pandemic. We also provide evidence for early community-based transmission and catalogue new mutations in SARS-CoV-2 strains in the UAE. Our findings contribute to the understanding of the global transmission network of SARS-CoV-2.

## Introduction

In December 2019, several cases of a new respiratory illness (now called COVID-19) were reported in the city of Wuhan (Hubei province, China) and in January 2020 it was confirmed these infections were caused by a novel coronavirus subsequently named SARS-CoV-2 [1–2]. On 12 March 2020, the ongoing SARS-CoV-2 outbreak was declared a pandemic by the World Health Organization (WHO) [3]. As of 14 August 2020, there have been 20.9 million laboratory-confirmed cases of COVID-19 and more than 760,000 deaths in 188 countries [4].

Dubai in the United Arab Emirates (UAE) is a cosmopolitan metropolis that has become a popular tourist destination and home to one of the busiest airport hubs in the world connecting the east with the west [2, 5]. Currently, the UAE has reported, 63,819 confirmed cases and 359 COVID-19-associated deaths (0.6% case fatality; 14 August 2020) [4]. In view of Dubai’s important tourism and travel connections, we attempted to characterize the full-genome sequence of SARS-CoV-2 strains from the index and early patients with COVID-19 in Dubai to gain a deeper understanding of the molecular epidemiology of the outbreak in Asia, Europe, and the Middle East.

## Results

### Patient Cohort and SARS-CoV-2 Whole Genome Sequencing

The 49 patients included in this study were the earliest confirmed cases in the UAE. The time period of 29 January to 18 March 2020 was specifically selected to focus on early SARS-CoV-2 viral introductions into the UAE. The first index patient in the UAE was reported on the 29 January 2020. Subsequently, Emirates airlines suspended flights to and from 30 global destinations from 18 March 2020 and Dubai airport was closed to passenger flights on 25 March 2020; hence, patients after 18 March 2020 were expected to be more likely a result of community transmission as opposed to imported infections. The index patient in the UAE was a female Chinese tourist (aged 63 years) travelling from Wuhan with other family members to visit her son in Dubai. The Chinese family arrived in Dubai on 16 January 2020 and tested positive on the 29 January 2020 (Table 1). Over the next seven weeks, there were multiple new cases among tourists and residents with travel history (44.9% had travel history from Europe) (Table 1). Nearly two-thirds (63.3%) of patients were male and 61.2% were aged between 20 and 44 years reflecting the young age structure of the UAE population [5]. Majority of patients (88%) were asymptomatic or had mild symptoms and only four required intensive care with invasive ventilation (one death; Table 1).

**Table 1.**
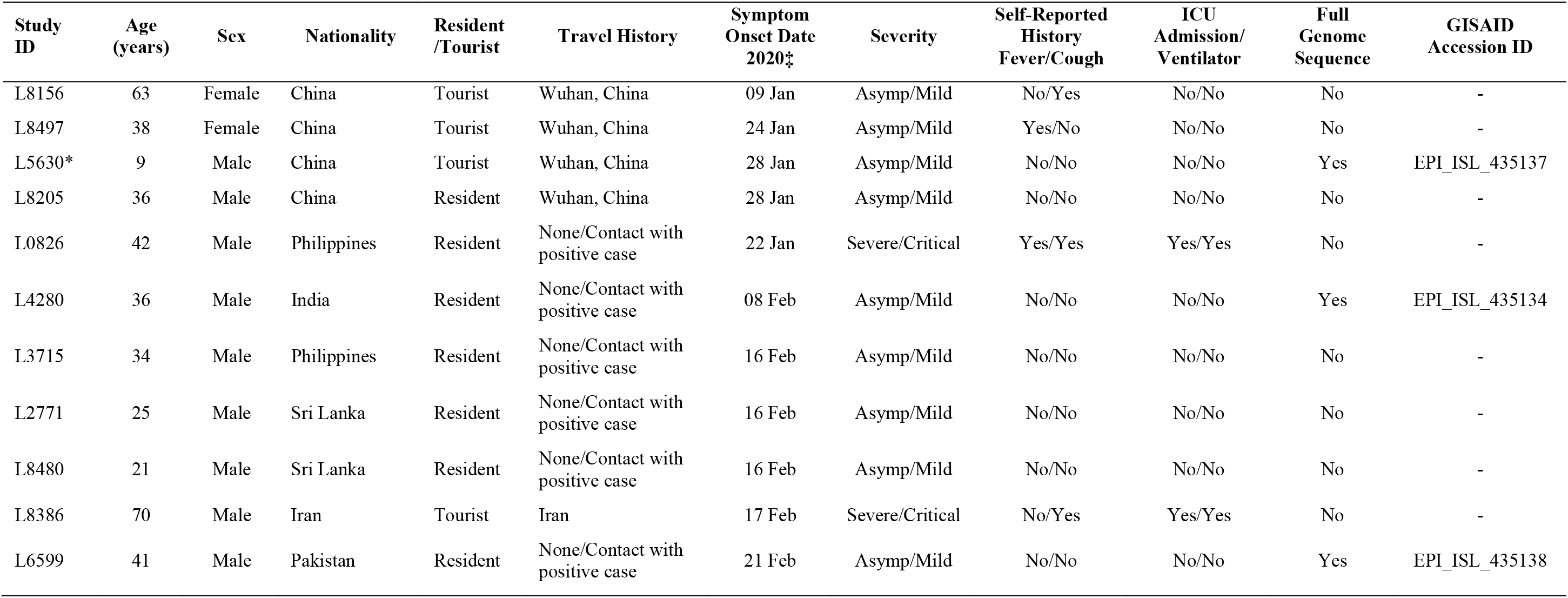
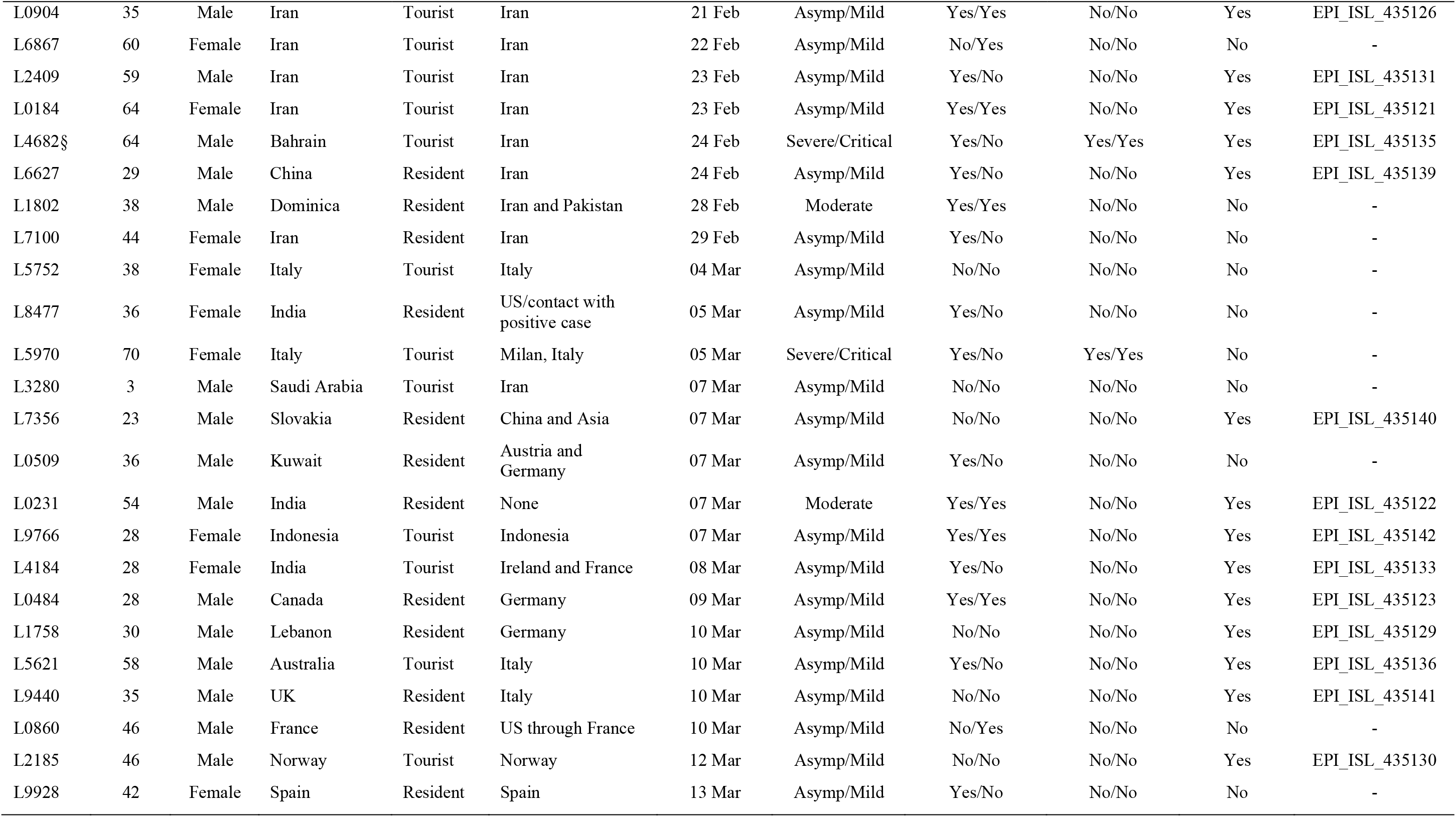
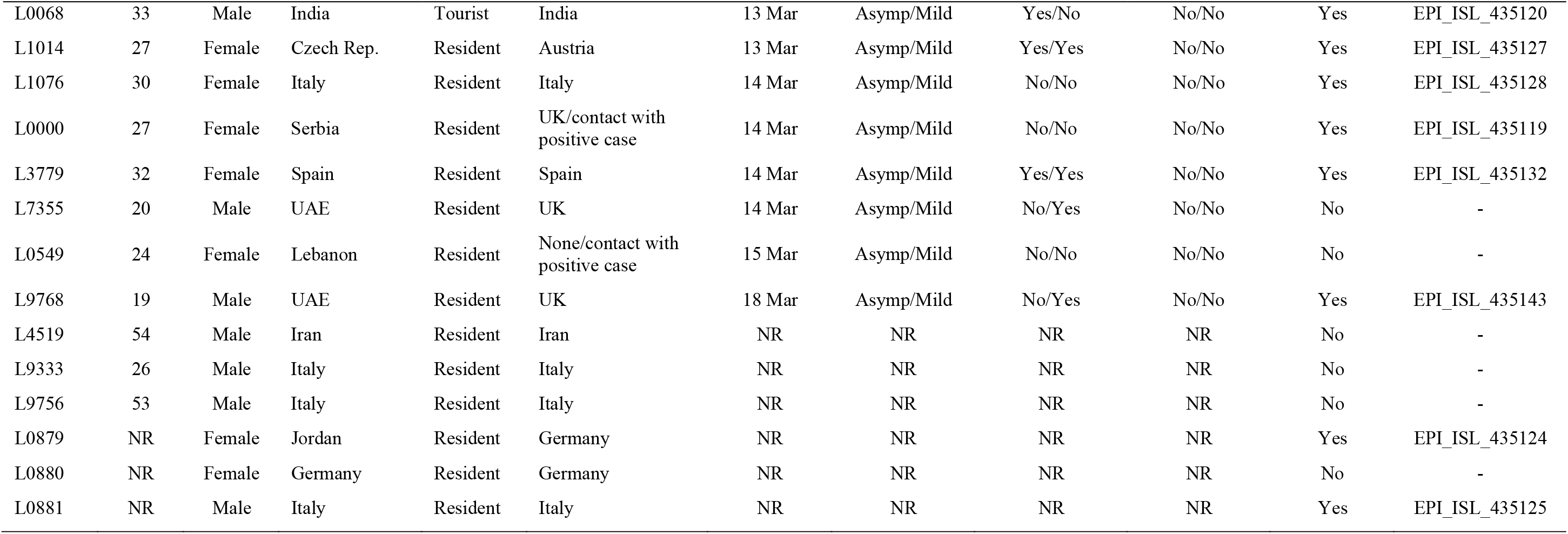
Sociodemographic and clinical characteristics of the index and early patients (n=49) with laboratory-confirmed SARS-CoV2 in Dubai, United Arab Emirates, 29 January-18 March 2020. *There was a second grandchild in the Chinese family cluster from Wuhan who remained negative throughout all testing. ‡Self-reported onset of symptoms extracted from medical records; NR, Not Reported; UK, United Kingdom; UAE, United Arab Emirates; US, United States. §deceased.

SARS-CoV-2 whole genome sequencing was performed on all 49 COVID-19 patient samples. Only genomes with almost complete coverage (n=25, Methods) were used for phylogenetic analysis. The 25 genomes were obtained from cases with disease onset in late January (n=1), early February (n=1), late February (n=6), early March (n=8), and late March (n=9). Of those, approximately two-thirds were male and aged between 10 and 40 years (Table 1).

### Phylogenetic Analysis

To understand early viral transmission in Dubai in the global context, we performed phylogenetic analysis on the 25 novel viral genomes we sequenced from early patients in the UAE (Table 1) in this study (Methods) along with 157 largely complete SARS-CoV-2 genomes deposited in GISAID from different countries between December 2019 and early March 2020 [6,7] (Table S1).

Consistent with multiple independent introductions, the UAE SARS-CoV-2 isolates were distributed across the phylogenetic tree (Figure 1). The majority (76%) clustered with clades A2a (48%) and A3 (28%) which are largely composed of isolates from COVID-19 patients in Europe and Iran, respectively. This clearly suggests that the major introductions into the UAE during the early phase of the pandemic originated from Europe and the Middle East/Iran.

**Figure 1.**
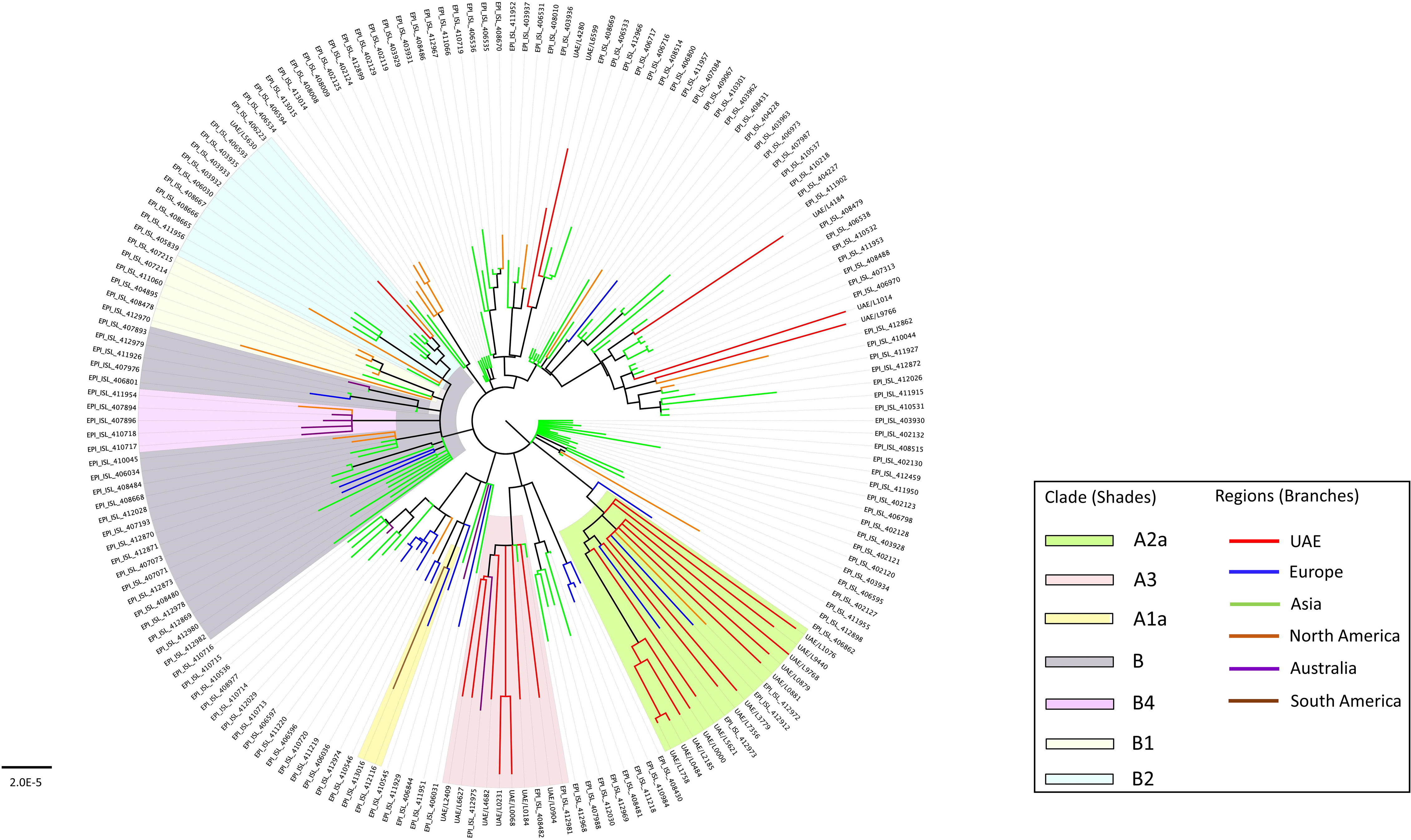
Phylogenetic relationships of SARS-CoV-2 isolates from early patients in Dubai and early global strains. A maximum likelihood phylogeny of 182 SARS-CoV-2 genomes (157 obtained from GISAID as of early Match and 25 genomes in this study). Bootstrap values >70% supporting major branches are shown. The 3 non-UAE isolates in clades A2a and A3 namely, Mexico/CDMX-InDRE_01, Germany/Baden/Wuerttemberg-1, and Australia/NSW05 are the GISAID ID: EPI_ISL_412972, GISAID ID: EPI_ISL_412912, and GISAID ID: EPI_ISL_412975, respectively, referred to in the main text. Scale bar represents number of nucleotide substitutions per site. UAE = United Arab Emirates.

Supporting its European origin, all individuals with the A2a clade isolates were mostly European and/or with recent travel history to a European country, mainly to Italy (n=4), Germany (n=3), United Kingdom (n=2), Spain (n=1), and Norway (n=1) (Table 1 and Figure 2). Onset of symptoms reported in this group was within or after the second week of March (Table 1) suggesting that the viral infections in this group could have occurred during late February to early March. Of note, a SARS-CoV-2 isolate submitted from Mexico (GISAID ID: EPI_ISL_412972) was 100% identical to that from an Italian expatriate working in the UAE (L0881), while another submitted in Germany (GISAID ID: EPI_ISL_412912) differed by a single mutation (Figure 1). All three individuals had a recent travel history to Italy and overlapping infection time frames (late February – early March). Within this group, isolates from patients L1758, L0484, and L2185 were identical (Figure 2) suggesting a possible common direct source of transmission.

**Figure 2.**
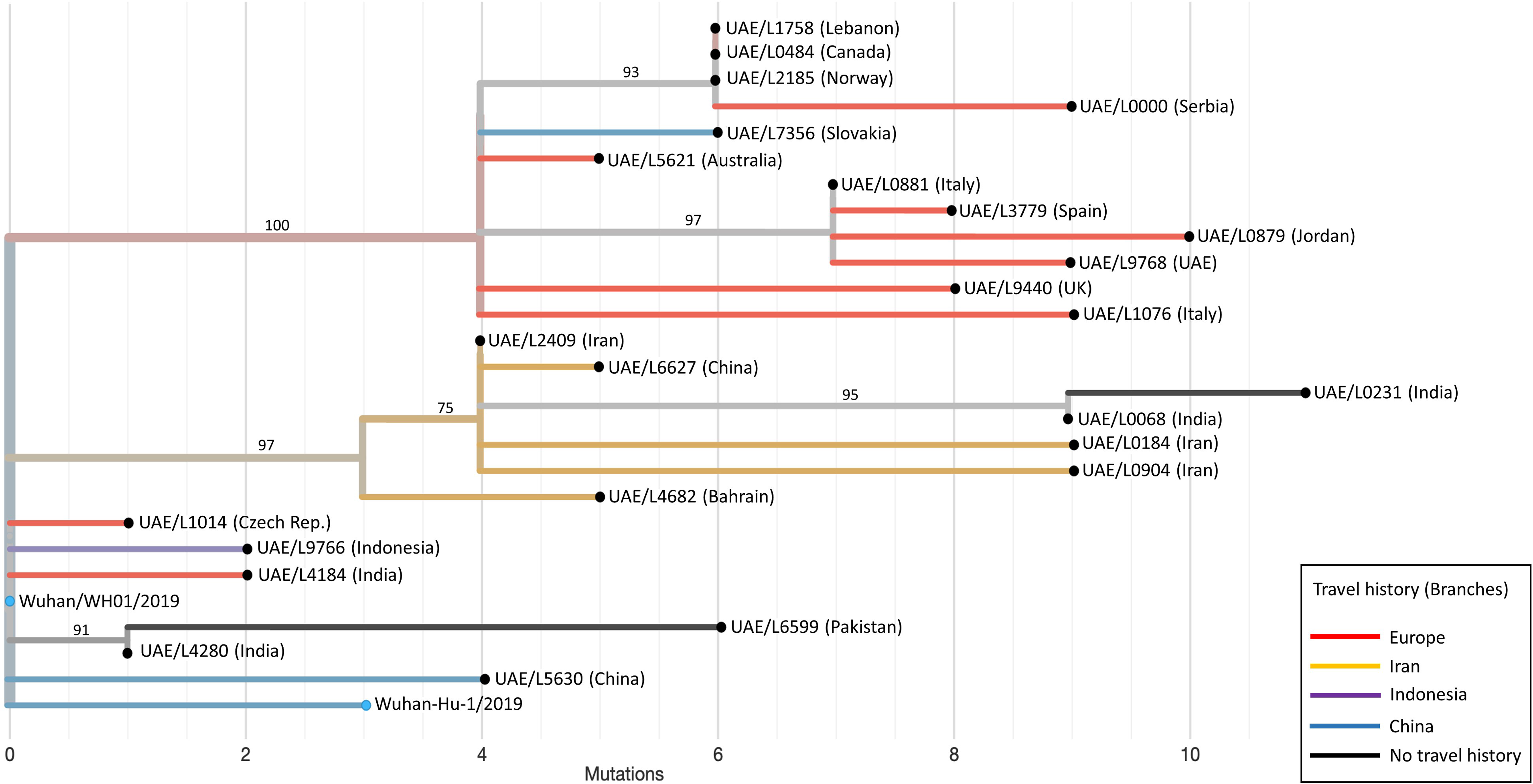
Relationship of early SARS-CoV-2 isolates in the UAE based on phylogenetic analysis and patients’ travel history. A maximum likelihood phylogenetic tree of all 25 UAE SARS-CoV-2 sequences, generated in this study, is shown. The two Wuhan genomes (Wuhan-Hu-1/2019, GISAID ID: EPI_ISL_402125 and Wuhan/WH01/2019, GISAID ID: EPI_ISL_406798) were used as reference genomes (blue filled circles). UAE viral strains (black filled circles) were labelled with sample ID and Nationality (in brackets). Bootstrap values >70% supporting major branches are shown. Branch lengths mark divergence from the reference Wuhan SARS-CoV-2 genome (GenBank accession number: NC_045512.2) in mutations numbers, while branch colour represents travel history. Travel history to Europe for patients UAE/L1758, UAE/L0484, UAE/L2185, and UAE/L0881 is obscured by vertical lines. UAE = United Arab Emirates.

Isolates in the A3 clade were obtained from five individuals with travel history to Iran (L2409, L6627, L0904, L0184, and L4682), one Indian resident (L0231), and one Indian tourist (L0068) (Figure 2). Onset of symptoms for the five individuals with travel history in this group was reported to be around 21-24 February (Table 1). Patient L0231 had no travel history and reported symptom onset on 7 March suggesting a possible community-based transmission event. Interestingly, all but one isolate obtained from patient L4682 – the only patient in this group with severe clinical presentation – shared a common ancestral strain identical to that obtained from patient L2409. The SARS-CoV-2 isolate from L4682 had two unique missense variants in the ORF1ab gene (Table S3) which might be worth investigating for any possible biological effect(s). Consistent with its Iranian origin, a SARS-CoV-2 sequence submitted by the University of Sydney (GISAID ID: EPI_ISL_412975) on 28 February 2020 differed by only two mutations from that of L2409, and both this Iranian male tourist and the Australian male had a recent travel history to Iran. We speculate that individuals with travel history to Iran around this time frame (L8386, L6867, and L3280), for whom a full viral genome sequence could not be obtained, were also very likely to cluster within the A3 clade.

Only one viral strain obtained from L5630, a family member of the early Chinese index patient, belonged to the B2 clade. Although we did not obtain full viral genome sequences from the other members of that Chinese family, we expect that all had a similar strain to L5630. Interestingly, our data do not suggest any transmission of this clade at least among the earliest patients (Figure 2) included in this study which is consistent with the reported early detection and isolation of this family. This finding also supports the notion of secondary source(s) for the ongoing local transmission.

The remaining five isolates did not belong to A2a, A3, B2, or any of the clades on nextrain.org as of 12 May 2020, suggesting earlier introduction(s). Those isolates were obtained from four Asians, two residents (L4280, L6599) and two tourists (L4184, L9766), and one Czech resident (L1014) working as an airline cabin crew with travel history to Austria (Table 1). Consistent with the Asian predominance among this patient group and the fewer (1 or 2) mutations for most of their isolates (4 out of 5) relative to the Wuhan reference genome (Figure 2), several early viral strains submitted in Asia clustered very closely to this group (Figure 1). L4280 was the first sequenced patient without travel history and became infected after transporting a work colleague, L0826, to hospital. Patient L0826 reported symptoms onset on 22 January suggesting that community-based transmission started in the UAE in early-to-mid January. L6599 was an Indian expatriate living with three other Filipino and Sri Lankan expatriates (L3715, L2771, L8480) (Table 1). All four individuals had no documented recent travel history suggesting local transmission, and although full viral genome sequences could only be obtained from one patient L6599, it is very likely that all have related isolates.

In aggregate, we identified 70 variants relative to the reference GenBank SARS-CoV-2 sequence NC_045512.2. The majority of these variants were missense (n=41) with the most frequent nucleotide change being C>T (n=33), and more than half (38/70) were localized in the ORF1ab gene (Table S3). Notably, 2 out of the 70 variants were novel as they were not identified in the Chinese National Center for Bioinformation Database (https://bigd.big.ac.cn/ncov/variation/annotation; last accessed August 13, 2020). The novel variants were a coding missense variant and a synonymous variant in the N and ORF1ab genes, respectively. In addition, 9 variants were very rare (i.e. seen less than 4 times out of 81,625 genomes), including one missense variant (F850I) in the S gene (Table S3).

## Discussion

Our findings suggest multiple independent spatiotemporal introductions of SARS-CoV-2 into the UAE where the majority of introductions (76%) were from Iran and Europe during two different time frames (mid-late February and early March, respectively). Although we show evidence for possible local transmission within the Middle Eastern/Iranian isolates, it will be important to sequence further isolates at subsequent dates to determine whether these introductions succeeded in seeding more clustering and whether such clustering was affected by proactive and vigilant public health measures, such as transitioning to online learning for schools and universities, implementing work-from-home protocols across all sectors, and nationwide disinfection campaigns.

Six isolates (22%) did not cluster with the European or Iranian groups and represented earlier introductions which did not appear to seed larger clusters in our sampled cohort. However, additional sequencing is needed to determine the extent of community transmission, especially given that our data strongly suggest that the earliest patient (early to mid-January) in the UAE could have been a secondary infection from one of those introductions.

The new SARS-CoV-2 mutations identified in the UAE warrant further investigation to explore whether they influence viral characteristics, especially pathogenicity, or provide important information for vaccine development. One of the major strengths of the study was the non-biased representative sample of early cases, including the index family cluster, in Dubai from the only central testing lab, along with detailed demographic and clinical information. Limitations included the inability to conduct full whole genome sequencing on more samples most likely due to low viral load issues, although we were able to deduce the origin of transmission in most of those individuals based on travel history. Regardless, this study contributes important molecular epidemiological data that can be used to further understand the global transmission network of SARS-CoV-2 [8].

## Methods

### Human Subjects and Ethics Approval

Sociodemographic and clinical data was extracted from the electronic medical records of the earliest 49 patients with laboratory confirmed SARS-CoV-2 from 29 January to 18 March 2020 using the WHO case report form. Cases were categorized into three groups based on disease severity: asymptomatic and mild cases with either no symptoms or mild non-life-threatening symptoms e.g. dry cough, mild fever; moderate cases with symptoms (e.g. breathlessness, persistent fever) requiring hospitalization and medical attention (e.g. supplementary oxygen therapy, intravenous fluids); and severe/critical cases with advanced disease and pneumonia requiring admission to intensive care units and specialized life-support treatment (e.g. mechanical ventilation). This study was approved by the Dubai Scientific Research Ethics Committee - Dubai Health Authority (approval number #DSREC-04/2020_02). The requirement for informed consent was waived as this study was part of a public health surveillance and outbreak investigation in the UAE. Nonetheless, all patients treated at a healthcare facility in the UAE provide written consent for their deidentified data to be used for research and this study was performed in accordance with the relevant laws and regulations that govern research in the UAE.

### SARS-CoV-2 Whole Genome Sequencing

All 49 COVID-19 patients tested positive for SARS-CoV-2 by RT-qPCR using RNA extracted from nasopharyngeal swabs following the QIAamp Viral RNA Mini or the EZ1 DSP Virus Kits (Qiagen, Hilden, Germany). RNA libraries from all samples were then prepared for shotgun transcriptomic sequencing using the TruSeq Stranded Total RNA Library kit from Illumina (San Diego, CA, USA), following manufacturer’s instructions. Libraries were sequenced using the NovaSeq SP Reagent kit (2 × 150 cycles) from Illumina (San Diego, CA, USA). Sample L5630 underwent a target enrichment approach where double stranded DNA (synthesized using the QuantiTect Reverse Transcription Kit from Qiagen, Hilden, Germany) was amplified using 26 overlapping primer sets covering most of the SARS-CoV-2 genome as recently described by our group [9]. PCR products were then sheared by ultra-sonication (Covaris LE220-plus series, MA, USA) and prepared for sequencing using the SureSelectXT Library Preparation kit (Agilent, CA, USA). This library was sequenced using the Illumina MiSeq Micro Reagent Kit, V2 (2 × 150 cycles).

### SARS-CoV-2 Genome Assembly

High quality (>Q30) sequencing reads were trimmed and then aligned to the reference SARS-CoV-2 genome from Wuhan, China (GenBank accession number: NC_045512.2) using a custom-made bioinformatics pipeline (Fig. S1). Assembled genomes with at least 20X average coverage across most nucleotide positions (56-29,797) were used for subsequent phylogenetic analysis (Table S1). A total of 25 viral genomes (24 by shotgun and 1 by target enrichment) met this inclusion criterion and were submitted to the Global Initiative on Sharing All Influenza Data (GISAID) database under accession IDs: EPI_ISL_435119-435142 (Table S2).

### Phylogenetic Analysis

We downloaded 157 global non-UAE sequences (Table S2) with largely complete genomes (nucleotide positions 56-29,797) submitted to GISAID EpiCoV (https://www.epicov.org/) between December 2019 and 04 March 2020 [7]. All 182 sequences, including the 25 UAE sequences generated in this study, were analysed using Nexstrain [10], which consists of Augur v6.4.3 pipeline for multiple sequence alignment (MAFFT v7.455 [11]) and phylogenetic tree construction (IQtree v1.6.12 [12]). Tree topology was assessed using the fast bootstrapping function with 1000 replicates. Tree visualization and annotations were performed in FigTree v1.4.4 [13] for Figure 1 and in auspice v2.13.0 tool [10] for Figure 2. SARS-CoV-2 clades annotations were performed in auspice v2.13.0 and cross-checked with nextstrain.org as of 12 May 2020.

## About the Author

Dr. Abou Tayoun is an Associate Professor in the Department of Genetics, Mohammed Bin Rashid University of Medicine and Health Sciences. He is also the director of Al Jalila Genomics Center at Al Jalila Children’s Specialty Hospital. His research interests are focused around clinical and population genomics in general, and specifically in the Middle East.

## Author Contributions

AAA, AT, NN, and TL conceived the study and drafted the protocol. All authors provided critical input into the protocol. AAA, AAK, and HK coordinated the ethical approval and sample retrieval. RV and ZD conducted the RT-qPCR analysis. AT, SR, and DH performed the whole genome sequencing and phylogenetic analysis. HK performed data extraction from the medical records and TL completed data analysis for the manuscript. AT and TL drafted the manuscript, AAA and NN refined it before AAK, AS, DH HA, HK, MU, QH, RH, RHA, RV, SR, and ZD, provided comments and feedback on the first draft. All authors read and approved the final version of the manuscript.

## Data availability statement

All data generated or analysed during this study are included in this published article (and its Supplementary Information files) and the sequences are available on the GISAID database under the corresponding accession numbers.

