## Supplementary Information for "Multiple early introductions of SARS-CoV-2 into a global travel hub in the Middle East" (from another version of this preprint: version 3, 2020-06-17)

**Genomic surveillance and phylogenetic analysis reveal multiple introductions of SARS-CoV-2 into a global travel hub in the Middle East**

Ahmad Abou Tayoun^1,2 ‡^, Tom Loney^2 ‡^, Hamda Khansaheb^3^, Sathishkumar Ramaswamy^1^, Divinlal Harilal^1^, Zulfa Omar Deesi^4^, Rupa Murthy Varghese^4^, Hanan Al Suwaidi^2^, Abdulmajeed Alkhajeh^3^, Laila Mohamed AlDabal^5^, Mohammed Uddin^2,6^, Rifat Hamoudi^7^, Rabih Halwani^7^, Abiola Senok^2^, Qutayba Hamid^8^, Norbert Nowotny^2,9*^, Alawi Alsheikh-Ali^2*^

Corresponding authors: Norbert Nowotny,; Alawi Alsheikh-Ali,.

^‡^ Joint first authorship.

**This file includes:**

Figures S1

Tables S1 to S3

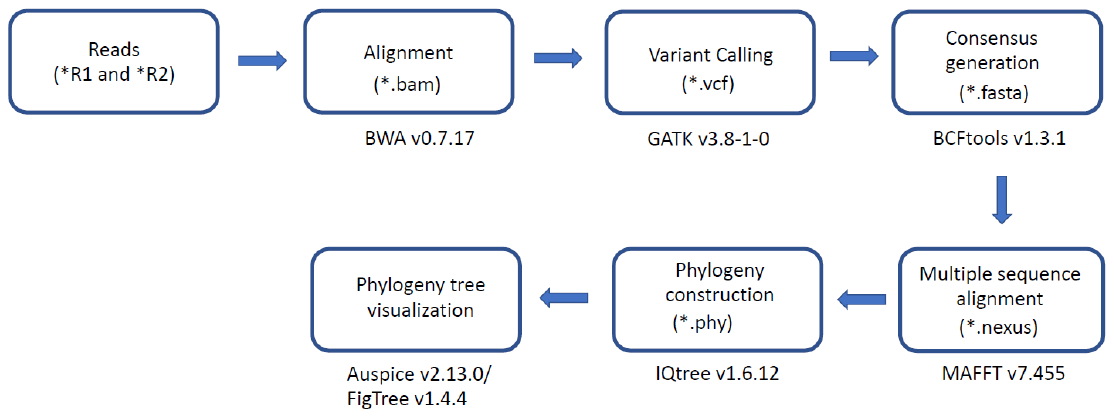

**Fig. S1**. Analysis pipeline starting with sequencing raw data to phylogenetic tree construction and visualization. Input and output file types for each programme are shown in brackets. The progamme name and version are indicated under each step. R1, sequencing read 1; R2, sequencing read 2.

Table S1. Sequencing and coverage statistics for all 25 UAE samples used for phylogenetic analysis in this study

| **Sample** | **Total Reads** | **Reads Aligned** | **% of Reads Aligned** | **Average Coverage*** | **% genome > 1X** | **% genome > 5X** | **% genome > 10X** | **% genome > 20X** |
| --- | --- | --- | --- | --- | --- | --- | --- | --- |
| UAE/L0000 | 33,700,572 | 208,432 | 0.62 | 43.23 | 100 | 100 | 100 | 97.3438 |
| UAE/L0068 | 35,660,576 | 261,236 | 0.73 | 125.57 | 100 | 100 | 100 | 100 |
| UAE/L0184 | 38,776,094 | 377,389 | 0.97 | 320.12 | 100 | 100 | 100 | 100 |
| UAE/L0231 | 27,406,298 | 451,672 | 1.65 | 367.38 | 100 | 100 | 100 | 100 |
| UAE/L0484 | 24,412,838 | 2,169,094 | 8.89 | 5252.05 | 100 | 100 | 100 | 100 |
| UAE/L0879 | 39,089,390 | 306,182 | 0.78 | 287.66 | 100 | 100 | 100 | 100 |
| UAE/L0881 | 33,480,022 | 419,908 | 1.25 | 153.05 | 100 | 100 | 100 | 100 |
| UAE/L0904 | 30,026,790 | 516,323 | 1.72 | 1028.81 | 100 | 100 | 100 | 100 |
| UAE/L1014 | 36,987,476 | 155,351 | 0.42 | 155.46 | 100 | 100 | 100 | 100 |
| UAE/L1076 | 36,202,168 | 191,552 | 0.53 | 182.22 | 100 | 100 | 100 | 100 |
| UAE/L1758 | 32,649,592 | 937,403 | 2.87 | 2106.37 | 100 | 100 | 100 | 100 |
| UAE/L2185 | 23,466,468 | 3,193,614 | 13.61 | 6492.55 | 100 | 100 | 100 | 100 |
| UAE/L2409 | 46,328,358 | 2,259,490 | 4.88 | 5499.47 | 100 | 100 | 100 | 100 |
| UAE/L3779 | 29,912,422 | 395,560 | 1.32 | 320.98 | 100 | 100 | 100 | 100 |
| UAE/L4184 | 25,150,950 | 215,797 | 0.86 | 31.14 | 100 | 99.93948 | 99.4318 | 85.697 |
| UAE/L4280 | 33,921,438 | 69,460 | 0.20 | 20.94 | 100 | 99.798265 | 97.2362 | 51.3348 |
| UAE/L4682 | 39,720,236 | 923,462 | 2.32 | 1877.56 | 100 | 100 | 100 | 100 |
| UAE/L5621 | 34,881,024 | 2,155,297 | 6.18 | 4679.84 | 100 | 100 | 100 | 100 |
| UAE/L5630 | 7,027,150 | 6,731,841 | 95.80 | 25339.89 | 100 | 100 | 100 | 100 |
| UAE/L6599 | 40,969,224 | 270,023 | 0.66 | 229.30 | 100 | 100 | 100 | 100 |
| UAE/L6627 | 37,417,892 | 566,560 | 1.51 | 988.90 | 100 | 100 | 100 | 100 |
| UAE/L7356 | 15,704,894 | 1,279,368 | 8.15 | 2523.43 | 100 | 100 | 100 | 100 |
| UAE/L9440 | 28,015,394 | 107,206 | 0.38 | 38.39 | 100 | 100 | 99.9731 | 95.6055 |
| UAE/L9766 | 30,981,012 | 731,611 | 2.36 | 1406.29 | 100 | 100 | 100 | 100 |
| UAE/L9768 | 22,370,982 | 150,635 | 0.67 | 91.97 | 100 | 99.989913 | 99.9798 | 99.9798 |

*Coverage calculated across positions 56-29797 (NC_045512.2).

Grey highlighted sample was sequenced using target enrichment approach (see Methods)

Table S2. Early UAE and non-UAE SARS-CoV-2 sequences analyzed in this study.

| **Non-UAE Strain** | **GISAID_epi_isl** | **GenBank accession** | **UAE Strain*** | **GISAID_epi_isl** |
| --- | --- | --- | --- | --- |
| BetaCov/Australia/NSW01/2020 | EPI_ISL_407893 |  | UAE/L0000 | [EPI_ISL_435119](https://www.epicov.org/epi3/start/EPI_ISL/435119) |
| BetaCov/Australia/NSW05/2020 | EPI_ISL_412975 |  | UAE/L0068 | [EPI_ISL_435120](https://www.epicov.org/epi3/start/EPI_ISL/435120) |
| BetaCov/Australia/QLD01/2020 | EPI_ISL_407894 |  | UAE/L0184 | [EPI_ISL_435121](https://www.epicov.org/epi3/start/EPI_ISL/435121) |
| BetaCov/Australia/QLD02/2020 | EPI_ISL_407896 |  | UAE/L0231 | [EPI_ISL_435122](https://www.epicov.org/epi3/start/EPI_ISL/435122) |
| BetaCov/Australia/QLD03/2020 | EPI_ISL_410717 |  | UAE/L0484 | [EPI_ISL_435123](https://www.epicov.org/epi3/start/EPI_ISL/435123) |
| BetaCov/Australia/QLD04/2020 | EPI_ISL_410718 |  | UAE/L0879 | [EPI_ISL_435124](https://www.epicov.org/epi3/start/EPI_ISL/435124) |
| BetaCov/Australia/VIC01/2020 | EPI_ISL_406844 | MT007544 | UAE/L0881 | [EPI_ISL_435125](https://www.epicov.org/epi3/start/EPI_ISL/435125) |
| BetaCov/Belgium/GHB-03021/2020 | EPI_ISL_407976 |  | UAE/L0904 | [EPI_ISL_435126](https://www.epicov.org/epi3/start/EPI_ISL/435126) |
| BetaCov/Brazil/SPBR-02/2020 | EPI_ISL_413016 | MT126808 | UAE/L1014 | [EPI_ISL_435127](https://www.epicov.org/epi3/start/EPI_ISL/435127) |
| BetaCov/Cambodia/0012/2020 | EPI_ISL_411902 |  | UAE/L1076 | [EPI_ISL_435128](https://www.epicov.org/epi3/start/EPI_ISL/435128) |
| BetaCov/Canada/ON-PHL2445/2020 | EPI_ISL_413014 |  | UAE/L1758 | [EPI_ISL_435129](https://www.epicov.org/epi3/start/EPI_ISL/435129) |
| BetaCov/Canada/ON/VIDO-01/2020 | EPI_ISL_413015 |  | UAE/L2185 | [EPI_ISL_435130](https://www.epicov.org/epi3/start/EPI_ISL/435130) |
| BetaCov/China/IQTC01/2020 | EPI_ISL_412966 |  | UAE/L2409 | [EPI_ISL_435131](https://www.epicov.org/epi3/start/EPI_ISL/435131) |
| BetaCov/China/IQTC02/2020 | EPI_ISL_412967 |  | UAE/L3779 | [EPI_ISL_435132](https://www.epicov.org/epi3/start/EPI_ISL/435132) |
| BetaCov/China/WH-09/2020 | EPI_ISL_411957 | MT093631 | UAE/L4184 | [EPI_ISL_435133](https://www.epicov.org/epi3/start/EPI_ISL/435133) |
| BetaCov/China/WHU01/2020 | EPI_ISL_406716 |  | UAE/L4280 | [EPI_ISL_435134](https://www.epicov.org/epi3/start/EPI_ISL/435134) |
| BetaCov/China/WHU02/2020 | EPI_ISL_406717 |  | UAE/L4682 | [EPI_ISL_435135](https://www.epicov.org/epi3/start/EPI_ISL/435135) |
| BetaCov/Chongqing/IVDC-CQ-001/2020 | EPI_ISL_408481 |  | UAE/L5621 | [EPI_ISL_435136](https://www.epicov.org/epi3/start/EPI_ISL/435136) |
| BetaCov/Chongqing/YC01/2020 | EPI_ISL_408478 |  | UAE/L5630 | [EPI_ISL_435137](https://www.epicov.org/epi3/start/EPI_ISL/435137) |
| BetaCov/Chongqing/ZX01/2020 | EPI_ISL_408479 |  | UAE/L6599 | [EPI_ISL_435138](https://www.epicov.org/epi3/start/EPI_ISL/435138) |
| BetaCov/England/01/2020 | EPI_ISL_407071 |  | UAE/L6627 | [EPI_ISL_435139](https://www.epicov.org/epi3/start/EPI_ISL/435139) |
| BetaCov/England/02/2020 | EPI_ISL_407073 |  | UAE/L7356 | [EPI_ISL_435140](https://www.epicov.org/epi3/start/EPI_ISL/435140) |
| BetaCov/England/09c/2020 | EPI_ISL_412116 |  | UAE/L9440 | [EPI_ISL_435141](https://www.epicov.org/epi3/start/EPI_ISL/435141) |
| BetaCov/Foshan/20SF207/2020 | EPI_ISL_406534 |  | UAE/L9766 | [EPI_ISL_435142](https://www.epicov.org/epi3/start/EPI_ISL/435142) |
| BetaCov/Foshan/20SF210/2020 | EPI_ISL_406535 |  | UAE/L9768 | [EPI_ISL_435143](https://www.epicov.org/epi3/start/EPI_ISL/435143) |
| BetaCov/Foshan/20SF211/2020 | EPI_ISL_406536 |  |  |  |
| BetaCov/France/IDF0372-isl/2020 | EPI_ISL_410720 |  |  |  |
| BetaCov/France/IDF0372/2020 | EPI_ISL_406596 |  |  |  |
| BetaCov/France/IDF0373/2020 | EPI_ISL_406597 |  |  |  |
| BetaCov/France/IDF0386-islP1/2020 | EPI_ISL_411219 |  |  |  |
| BetaCov/France/IDF0386-islP3/2020 | EPI_ISL_411220 |  |  |  |
| BetaCov/France/IDF0515-isl/2020 | EPI_ISL_410984 |  |  |  |
| BetaCov/France/IDF0515/2020 | EPI_ISL_408430 |  |  |  |
| BetaCov/France/IDF0571/2020 | EPI_ISL_411218 |  |  |  |
| BetaCov/France/IDF0626/2020 | EPI_ISL_408431 |  |  |  |
| BetaCov/Fujian/13/2020 | EPI_ISL_411066 |  |  |  |
| BetaCov/Fujian/8/2020 | EPI_ISL_411060 |  |  |  |
| BetaCov/Germany/Baden-Wuerttemberg-1/2020 | EPI_ISL_412912 |  |  |  |
| BetaCov/Germany/BavPat1/2020 | EPI_ISL_406862 |  |  |  |
| BetaCov/Guangdong/20SF012/2020 | EPI_ISL_403932 |  |  |  |
| BetaCov/Guangdong/20SF013/2020 | EPI_ISL_403933 |  |  |  |
| BetaCov/Guangdong/20SF014/2020 | EPI_ISL_403934 |  |  |  |
| BetaCov/Guangdong/20SF025/2020 | EPI_ISL_403935 |  |  |  |
| BetaCov/Guangdong/20SF028/2020 | EPI_ISL_403936 |  |  |  |
| BetaCov/Guangdong/20SF040/2020 | EPI_ISL_403937 |  |  |  |
| BetaCov/Guangdong/20SF174/2020 | EPI_ISL_406531 |  |  |  |
| BetaCov/Guangdong/20SF201/2020 | EPI_ISL_406538 |  |  |  |
| BetaCov/Guangzhou/20SF206/2020 | EPI_ISL_406533 |  |  |  |
| BetaCov/Hangzhou/HZ-1/2020 | EPI_ISL_406970 | MT039873 |  |  |
| BetaCov/Hangzhou/HZCDC0001/2020 | EPI_ISL_407313 |  |  |  |
| BetaCov/Hefei/2/2020 | EPI_ISL_412026 |  |  |  |
| BetaCov/Hong Kong/VB20026565/2020 | EPI_ISL_412030 |  |  |  |
| BetaCov/Hong Kong/VM20001061/2020 | EPI_ISL_412028 |  |  |  |
| BetaCov/Hong Kong/VM20001988/2020 | EPI_ISL_412029 |  |  |  |
| BetaCov/Italy/CDG1/2020 | EPI_ISL_412973 |  |  |  |
| BetaCov/Italy/INMI1-cs/2020 | EPI_ISL_410546 |  |  |  |
| BetaCov/Italy/INMI1-isl/2020 | EPI_ISL_410545 |  |  |  |
| BetaCov/Italy/SPL1/2020 | EPI_ISL_412974 |  |  |  |
| BetaCov/Japan/AI/I-004/2020 | EPI_ISL_407084 |  |  |  |
| BetaCov/Japan/Hu_DP_Kng_19-020/2020 | EPI_ISL_412968 |  |  |  |
| BetaCov/Japan/Hu_DP_Kng_19-027/2020 | EPI_ISL_412969 |  |  |  |
| BetaCov/Japan/KY-V-029/2020 | EPI_ISL_408669 | LC522972 |  |  |
| BetaCov/Japan/NA-20-05-1/2020 | EPI_ISL_410531 |  |  |  |
| BetaCov/Japan/OS-20-07-1/2020 | EPI_ISL_410532 |  |  |  |
| BetaCov/Japan/TY-WK-012/2020 | EPI_ISL_408665 | LC522973 |  |  |
| BetaCov/Japan/TY-WK-501/2020 | EPI_ISL_408666 | LC522974 |  |  |
| BetaCov/Japan/TY-WK-521/2020 | EPI_ISL_408667 | LC522975 |  |  |
| BetaCov/Jiangsu/IVDC-JS-001/2020 | EPI_ISL_408488 |  |  |  |
| BetaCov/Jiangsu/JS01/2020 | EPI_ISL_411950 |  |  |  |
| BetaCov/Jiangsu/JS02/2020 | EPI_ISL_411952 |  |  |  |
| BetaCov/Jiangsu/JS03/2020 | EPI_ISL_411953 |  |  |  |
| BetaCov/Jiangxi/IVDC-JX-002/2020 | EPI_ISL_408486 |  |  |  |
| BetaCov/Jingzhou/HBCDC-HB-01/2020 | EPI_ISL_412459 |  |  |  |
| BetaCov/Korea/KCDC05/2020 | EPI_ISL_412869 |  |  |  |
| BetaCov/Korea/KCDC06/2020 | EPI_ISL_412870 |  |  |  |
| BetaCov/Korea/KCDC07/2020 | EPI_ISL_412871 |  |  |  |
| BetaCov/Korea/KCDC12/2020 | EPI_ISL_412872 |  |  |  |
| BetaCov/Korea/KCDC24/2020 | EPI_ISL_412873 |  |  |  |
| BetaCov/Mexico/CDMX/InDRE_01/2020 | EPI_ISL_412972 |  |  |  |
| BetaCov/Nepal/61/2020 | EPI_ISL_410301 | MT072688 |  |  |
| BetaCov/Nonthaburi/61/2020 | EPI_ISL_403962 |  |  |  |
| BetaCov/Nonthaburi/74/2020 | EPI_ISL_403963 |  |  |  |
| BetaCov/Shandong/IVDC-SD-001/2020 | EPI_ISL_408482 |  |  |  |
| BetaCov/Shenzhen/HKU-SZ-002/2020 | EPI_ISL_406030 | MN938384 |  |  |
| BetaCov/Shenzhen/HKU-SZ-005/2020 | EPI_ISL_405839 | MN975262 |  |  |
| BetaCov/Shenzhen/SZTH-002/2020 | EPI_ISL_406593 |  |  |  |
| BetaCov/Shenzhen/SZTH-003/2020 | EPI_ISL_406594 |  |  |  |
| BetaCov/Shenzhen/SZTH-004/2020 | EPI_ISL_406595 |  |  |  |
| BetaCov/Sichuan/IVDC-SC-001/2020 | EPI_ISL_408484 |  |  |  |
| BetaCov/Singapore/1/2020 | EPI_ISL_406973 |  |  |  |
| BetaCov/Singapore/10/2020 | EPI_ISL_410716 |  |  |  |
| BetaCov/Singapore/11/2020 | EPI_ISL_410719 |  |  |  |
| BetaCov/Singapore/2/2020 | EPI_ISL_407987 |  |  |  |
| BetaCov/Singapore/3/2020 | EPI_ISL_407988 |  |  |  |
| BetaCov/Singapore/5/2020 | EPI_ISL_410536 |  |  |  |
| BetaCov/Singapore/6/2020 | EPI_ISL_410537 |  |  |  |
| BetaCov/Singapore/7/2020 | EPI_ISL_410713 |  |  |  |
| BetaCov/Singapore/8/2020 | EPI_ISL_410714 |  |  |  |
| BetaCov/Singapore/9/2020 | EPI_ISL_410715 |  |  |  |
| BetaCov/South Korea/KCDC03/2020 | EPI_ISL_407193 |  |  |  |
| BetaCov/South Korea/SNU01/2020 | EPI_ISL_411929 | MT039890 |  |  |
| BetaCov/Sweden/01/2020 | EPI_ISL_411951 | MT093571 |  |  |
| BetaCov/Sydney/3/2020 | EPI_ISL_408977 |  |  |  |
| BetaCov/Taiwan/2/2020 | EPI_ISL_406031 |  |  |  |
| BetaCov/Taiwan/3/2020 | EPI_ISL_411926 |  |  |  |
| BetaCov/Taiwan/4/2020 | EPI_ISL_411927 |  |  |  |
| BetaCov/Taiwan/CGMH-CGU-01/2020 | EPI_ISL_411915 |  |  |  |
| BetaCov/Taiwan/NTU02/2020 | EPI_ISL_410218 | MT066176 |  |  |
| BetaCov/USA/AZ1/2020 | EPI_ISL_406223 | MN997409 |  |  |
| BetaCov/USA/CA1/2020 | EPI_ISL_406034 | MN994467 |  |  |
| BetaCov/USA/CA2/2020 | EPI_ISL_406036 | MN994468 |  |  |
| BetaCov/USA/CA3/2020 | EPI_ISL_408008 | MT027062 |  |  |
| BetaCov/USA/CA4/2020 | EPI_ISL_408009 | MT027063 |  |  |
| BetaCov/USA/CA5/2020 | EPI_ISL_408010 | MT027064 |  |  |
| BetaCov/USA/CA6/2020 | EPI_ISL_410044 | MT044258 |  |  |
| BetaCov/USA/CA7/2020 | EPI_ISL_411954 | MT106052 |  |  |
| BetaCov/USA/CA8/2020 | EPI_ISL_411955 | MT106053 |  |  |
| BetaCov/USA/CA9/2020 | EPI_ISL_412862 | MT118835 |  |  |
| BetaCov/USA/IL2/2020 | EPI_ISL_410045 | MT044257 |  |  |
| BetaCov/USA/MA1/2020 | EPI_ISL_409067 | MT039888 |  |  |
| BetaCov/USA/TX1/2020 | EPI_ISL_411956 | MT106054 |  |  |
| BetaCov/USA/WA1-A12/2020 | EPI_ISL_407214 | MT020880 |  |  |
| BetaCov/USA/WA1-F6/2020 | EPI_ISL_407215 | MT020881 |  |  |
| BetaCov/USA/WA1/2020 | EPI_ISL_404895 | MN985325 |  |  |
| BetaCov/USA/WA2/2020 | EPI_ISL_412970 |  |  |  |
| BetaCov/USA/WI1/2020 | EPI_ISL_408670 | MT039887 |  |  |
| BetaCov/Vietnam/VR03-38142/2020 | EPI_ISL_408668 |  |  |  |
| BetaCov/Wuhan-Hu-1/2019 | EPI_ISL_402125 | MN908947 |  |  |
| BetaCov/Wuhan/HBCDC-HB-01/2019 | EPI_ISL_402132 |  |  |  |
| BetaCov/Wuhan/HBCDC-HB-02/2019 | EPI_ISL_412898 |  |  |  |
| BetaCov/Wuhan/HBCDC-HB-02/2020 | EPI_ISL_412978 |  |  |  |
| BetaCov/Wuhan/HBCDC-HB-03/2019 | EPI_ISL_412899 |  |  |  |
| BetaCov/Wuhan/HBCDC-HB-03/2020 | EPI_ISL_412979 |  |  |  |
| BetaCov/Wuhan/HBCDC-HB-04/2020 | EPI_ISL_412980 |  |  |  |
| BetaCov/Wuhan/HBCDC-HB-05/2020 | EPI_ISL_412981 |  |  |  |
| BetaCov/Wuhan/HBCDC-HB-06/2020 | EPI_ISL_412982 |  |  |  |
| BetaCov/Wuhan/IPBCAMS-WH-01/2019 | EPI_ISL_402123 | MT019529 |  |  |
| BetaCov/Wuhan/IPBCAMS-WH-02/2019 | EPI_ISL_403931 | MT019530 |  |  |
| BetaCov/Wuhan/IPBCAMS-WH-03/2019 | EPI_ISL_403930 | MT019531 |  |  |
| BetaCov/Wuhan/IPBCAMS-WH-04/2019 | EPI_ISL_403929 | MT019532 |  |  |
| BetaCov/Wuhan/IPBCAMS-WH-05/2020 | EPI_ISL_403928 | MT019533 |  |  |
| BetaCov/Wuhan/IVDC-HB-01/2019 | EPI_ISL_402119 |  |  |  |
| BetaCov/Wuhan/IVDC-HB-04/2020 | EPI_ISL_402120 |  |  |  |
| BetaCov/Wuhan/IVDC-HB-05/2019 | EPI_ISL_402121 |  |  |  |
| BetaCov/Wuhan/IVDC-HB-envF13-20/2020 | EPI_ISL_408514 |  |  |  |
| BetaCov/Wuhan/IVDC-HB-envF13-21/2020 | EPI_ISL_408515 |  |  |  |
| BetaCov/Wuhan/WH01/2019 | EPI_ISL_406798 | LR757998 |  |  |
| BetaCov/Wuhan/WH03/2020 | EPI_ISL_406800 | LR757996 |  |  |
| BetaCov/Wuhan/WH04/2020 | EPI_ISL_406801 | LR757995 |  |  |
| BetaCov/Wuhan/WIV02/2019 | EPI_ISL_402127 | MN996527 |  |  |
| BetaCov/Wuhan/WIV04/2019 | EPI_ISL_402124 | MN996528 |  |  |
| BetaCov/Wuhan/WIV05/2019 | EPI_ISL_402128 | MN996529 |  |  |
| BetaCov/Wuhan/WIV06/2019 | EPI_ISL_402129 | MN996530 |  |  |
| BetaCov/Wuhan/WIV07/2019 | EPI_ISL_402130 | MN996531 |  |  |
| BetaCov/Yu--an/IVDC-YN-003/2020 | EPI_ISL_408480 | MT049951 |  |  |
| BetaCov/Zhejiang/WZ-01/2020 | EPI_ISL_404227 |  |  |  |
| BetaCov/Zhejiang/WZ-02/2020 | EPI_ISL_404228 |  |  |  |

*Sequences generated in this study.

Table S3. SARS-CoV-2 sequence variants identified in this study.

| **Genomic position*** | **Nucleotide change** | **Gene name/Region** | **Amino acid change** | **Protein effect** | **Virus number with variation**** | **Samples** |
| --- | --- | --- | --- | --- | --- | --- |
| 61 | G>T | *5'UTR* |  | Noncoding | 3 | UAE/L0904 |
| 241 | C>T | *5'UTR* |  | Noncoding | 4676 | UAE/L5621,UAE/L9440,UAE/L1758,UAE/L7356,UAE/L0000,UAE/L0484,UAE/L2185,UAE/L1076,UAE/L3779,UAE/L0879,UAE/L0881,UAE/L9768 |
| 313 | C>T | *ORF1ab* | L16= | Synonymous | 62 | UAE/L3779 |
| 514 | TGTTATG>T | *ORF1ab* | 84_85del | Inframe deletion | 7 | UAE/L9768 |
| 520 | G>T | *ORF1ab* | M85I | Missense | 0 | UAE/L9766 |
| 884 | C>T | *ORF1ab* | R207C | Missense | 29 | UAE/L0068,UAE/L0231 |
| 1059 | C>T | *ORF1ab* | T265I | Missense | 1805 | UAE/L1758,UAE/L0000UAE/L0484,UAE/L2185 |
| 1170 | C>T | *ORF1ab* | S302F | Missense | 0 | UAE/L0184 |
| 1185 | C>T | *ORF1ab* | A307V | Missense | 0 | UAE/L0184 |
| 1397 | G>A | *ORF1ab* | V378I | Missense | 117 | UAE/L0904,UAE/L2409,UAE/L4682,UAE/L6627,UAE/L0184,UAE/L0068,UAE/L0231 |
| 1997 | C>T | *ORF1ab* | L578= | Synonymous | 3 | UAE/L9440 |
| 2246 | G>A | *ORF1ab* | G661S | Missense | 0 | UAE/L4682 |
| 2494 | G>A | *ORF1ab* | E743= | Synonymous | 0 | UAE/L4184 |
| 2724 | C>T | *ORF1ab* | P820L | Missense | 1 | UAE/L6627 |
| 3037 | C>T | *ORF1ab* | F924= | Synonymous | 4729 | UAE/L5621,UAE/L9440,UAE/L1758,UAE/L7356,UAE/L0000,UAE/L0484,UAE/L2185,UAE/L1076,UAE/L3779,UAE/L0879,UAE/L0881,UAE/L9768 |
| 3373 | C>A | *ORF1ab* | D1036E | Missense | 28 | UAE/L0879 |
| 4201 | G>T | *ORF1ab* | M1312I | Missense | 6 | UAE/L7356 |
| 4255 | G>T | *ORF1ab* | P1330= | Synonymous | 31 | UAE/L1076 |
| 8290 | C>T | *ORF1ab* | L2675= | Synonymous | 3 | UAE/L9440 |
| 8653 | G>T | *ORF1ab* | M2796I | Missense | 27 | UAE/L0068,UAE/L0231,UAE/1L076 |
| 8782 | C>T | *ORF1ab* | S2839= | Synonymous | 1278 | UAE/L5630 |
| 9246 | C>T | *ORF1ab* | A2994V | Missense | 5 | UAE/L4682 |
| 11074 | C>CTTT | *ORF1ab* | F3603dup | Inframe insertion | 9 | UAE/L0068 |
| 11083 | G>T | *ORF1ab* | L3603F | Missense | 863 | UAE/L0904,UAE/L2409,UAE/L6627,UAE/L0184,UAE/L0068 |
| 11195 | C>T | *ORF1ab* | L3644F | Missense | 9 | UAE/L0879 |
| 12880 | C>T | *ORF1ab* | I4205= | Synonymous | 17 | UAE/L5630 |
| 13568 | C>T | *ORF1ab* | A4435V | Missense | 18 | UAE/L5621 |
| 13957 | C>T | *ORF1ab* | R4565C | Missense | 0 | UAE/L6599 |
| 14408 | C>T | *ORF1ab* | P4715L | Missense | 4707 | UAE/L5621,UAE/L9440,UAE/L1758,UAE/L7356,UAE/L0000,UAE/L0484,UAE/L2185,UAE/L1076,UAE/L3779,UAE/L0879,UAE/L0881,UAE/L9768 |
| 14636 | C>T | *ORF1ab* | A4791V | Missense | 0 | UAE/L9766 |
| 15324 | C>T | *ORF1ab* | N5020= | Synonymous | 234 | UAE/L6599 |
| 16323 | C>T | *ORF1ab* | C5353= | Synonymous | 0 | UAE/L9768 |
| 16545 | T>C | *ORF1ab* | V5427= | Synonymous | 0 | UAE/L0184 |
| 17678 | C>T | *ORF1ab* | T5805M | Missense | 0 | UAE/L9440 |
| 18163 | A>G | *ORF1ab* | I5967V | Missense | 0 | UAE/L1014 |
| 18377 | C>T | *ORF1ab* | T6038I | Missense | 2 | UAE/L0184 |
| 18501 | C>T | *ORF1ab* | Y6079= | Synonymous | 2 | UAE/L4184 |
| 19073 | A>G | *ORF1ab* | D6270G | Missense | 1 | UAE/L0068,UAE/L0231 |
| 20235 | C>A | *ORF1ab* | P6657= | Synonymous | 2 | UAE/L0904 |
| 20787 | T>C | *ORF1ab* | N6841= | Synonymous | 1 | UAE/L0904 |
| 21644 | T>C | *S* | Y28H | Missense | 1 | UAE/L6599 |
| 21781 | C>T | *S* | T73= | Synonymous | 1 | UAE/L0000 |
| 22042 | T>C | *S* | Y160= | Synonymous | 1 | UAE/L0904 |
| 23170 | C>T | *S* | N536= | Synonymous | 0 | UAE/L6599 |
| 23403 | A>G | *S* | D614G | Missense | 4733 | UAE/L5621,UAE/L9440,UAE/L1758,UAE/L7356,UAE/L0000,UAE/L0484,UAE/L2185,UAE/L1076,UAE/L3779,UAE/L0879,UAE/L0881,UAE/L9768 |
| 24110 | A>T | *S* | F850I | Missense | 0 | UAE/L9440 |
| 24378 | C>T | *S* | S939F | Missense | 4 | UAE/L6599 |
| 25563 | G>T | *ORF3a* | Q57H | Missense | 2204 | UAE/L1758,UAE/L0000,UAE/L0484,UAE/L32185 |
| 26507 | A>T | *M* |  | Intergenic upstream variant | 0 | UAE/L1076 |
| 26527 | C>T | *M* | A2V | Missense | 6 | UAE/L7356 |
| 26530 | A>G | *M* | D3G | Missense | 82 | UAE/L1076 |
| 26775 | G>T | *M* | A85S | Missense | 1 | UAE/L0000 |
| 27746 | G>T | *ORF7a* | R118I | Missense | 0 | UAE/L1076 |
| 27915 | G>T | *ORF8* | G88 | Nonsense | 1 | UAE/L0068,UAE/L0231 |
| 28144 | T>C | *ORF8* | L84S | Missense | 1260 | UAE/L5630 |
| 28580 | G>T | *N* | D103Y | Missense | 21 | UAE/L0879 |
| 28688 | T>C | *N* | L139= | Synonymous | 115 | UAE/L0904,UAE/L2409,UAE/L4682,UAE/L6627,UAE/L0184,UAE/L0068,UAE/L0231 |
| 28770 | C>T | *N* | T166I | Missense | 0 | UAE/L9768 |
| 28835 | T>C | *N* | S188P | Missense | 9 | UAE/L0904 |
| 28868 | C>T | *N* | P199S | Missense | 1 | UAE/L0231 |
| 28881 | G>A | *N* | R203K | Missense | 1098 | UAE/L3779,UAE/L0879,UAE/L0881,UAE/L9768 |
| 28882 | G>A | *N* | R203= | Synonymous | 1096 | UAE/L3779,UAE/L0879,UAE/L0881,UAE/L9768 |
| 28883 | G>C | *N* | G204R | Missense | 1096 | UAE/L3779,UAE/L0879,UAE/L0881,UAE/L9768 |
| 29060 | A>G | *N* | T263A | Missense | 0 | UAE/L0904 |
| 29077 | C>T | *N* | Y268= | Synonymous | 2 | UAE/L0068,UAE/L0231 |
| 29095 | C>T | *N* | F274= | Synonymous | 33 | UAE/L5630 |
| 29303 | C>T | *N* | P344S | Missense | 11 | UAE/L6599,UAE/L4280 |
| 29374 | G>A | *N* | E367= | Synonymous | 12 | UAE/L0184 |
| 29415 | C>T | *N* | A381V | Missense | 2 | UAE/L0000 |
| 29742 | G>T | *3'UTR* |  | Noncoding | 165 | UAE/L0904,UAE/L2409,UAE/L4682,UAE/L6627,UAE/L0184,UAE/L0068,UAE/L0231 |

*Based on GenBank NC_045512.2 nucleotide reference sequence.

**Obtained from China National Center for Bioinformation (https://bigd.big.ac.cn/ncov/variation/annotation). Last accessed April 29, 2020.
